# Acute effects of systemic inflammation upon neurovascular unit and cerebrovascular function

**DOI:** 10.1101/498089

**Authors:** Gaia Brezzo, Julie Simpson, Kamar E. Ameen-Ali, Jason Berwick, Chris Martin

## Abstract

**Background:** Brain health relies on a tightly regulated system known as neurovascular function whereby the cellular constituents of the neurovascular unit (NVU) regulate cerebral haemodynamics in accordance with neuronal metabolic demand. Disruption of neurovascular function impairs brain health and is associated with the development of disease, including Alzheimer’s disease (AD). The NVU is the site of action of neuroinflammatory responses and contributes to the transition of systemic inflammation to neuroinflammatory processes. Thus, systemic inflammatory challenges may cause a shift in the NVU focus, prioritising neuroimmune over neurovascular actions leading to altered neurovascular function.

**Methods:** Rats were injected with lipopolysaccharide (LPS) (2mg/kg) or vehicle and haemodynamic responses to sensory and non-sensory (hypercapnia) stimuli were assessed *in vivo*. Following imaging, animals were perfused and their brain extracted to histologically characterise components of the NVU to determine the association between underlying pathology to altered blood flow regulation *in vivo*.

**Results:** LPS-treated animals showed altered haemodynamic function and cerebrovascular dynamics 6 hours after LPS administration. Histological assessment identified a significant increase in astrogliosis, microgliosis and endothelial activation in LPS-treated animals.

**Conclusions:** Our data shows that an acutely induced systemic inflammatory response is able to rapidly alter *in-vivo* haemodynamic function and is associated with significant changes in the cellular constituents of the NVU. We suggest that these effects are initially mediated by endothelial cells, which are directly exposed to the circulating inflammatory stimulus and have been implicated in regulating functional hyperaemia.

## 1.1 Background

Brain health and function are dependent on an extensively regulated blood supply. The local regulation of blood flow in accordance with metabolic neuronal demands in the tissue is termed neurovascular coupling and is orchestrated by the resident cells of the neurovascular unit (NVU). Disruption to neurovascular coupling impairs the delivery of critical substrates to brain cells and impedes the removal of by-products accumulated during cerebral metabolism [1]. Alterations of brain microenvironment and cellular interactions of the NVU have been implicated in the development of a number of neurodegenerative diseases, including Alzheimer’s disease (AD) [2-4]. Nevertheless the process or processes by which neurovascular function affects and is affected by neurodegenerative diseases *in vivo*, as well as the cellular substrates of these effects, remain unclear.

Neurovascular coupling underpins the physiological basis of non-invasive functional neuroimaging techniques, including functional magnetic resonance imaging (fMRI) and positron emission tomography (PET) in which changes to brain blood flow and oxygenation are tracked as surrogate markers for neuronal activity. Such neuroimaging techniques may provide new opportunities to predict, detect, diagnose and study brain disease processes using non-invasive imaging biomarkers, but are dependent on the mapping of relevant *in vivo* measurements to neuropathological changes in the context of specific disease processes such as inflammation.

Mounting evidence highlights inflammation as a major factor in the development of many neurodegenerative diseases, including AD [5-8]. Further evidence pinpoints inflammation as a driver of neuropathology [9] and it has been shown to precede the development of amyloid-beta (Aβ) plaques [10]. The NVU is the site of action of neuroinflammatory responses and contributes to the transition of systemic inflammation to neuroinflammatory processes. Several non-neuronal cells within the NVU are key players in the initiation and regulation of brain inflammatory responses, as well as in mediating the effects of systemic inflammation upon brain function. Activated astrocytes and microglia release a range of pro-inflammatory molecules [11-14]. Endothelial cells (ECs) also play an important role through upregulation of intercellular adhesion molecules (ICAM-1) and vascular cellular adhesion molecules (VCAM-1) [15].

Research has also highlighted a beneficial role for inflammation, suggesting that directing the inflammatory response may be of more therapeutic benefit than suppressing it [16, 17]. Glial cells have been shown to have a neuroprotective role in the neuroinflammatory response [18-21], highlighting the complexity and difficulty in pinpointing the roles and factors involved in the pathophysiological cascade of inflammation.

To help elucidate the changes that occur in the context of inflammation, numerous models have been developed. The peripheral lipopolysaccharide (LPS) injection method is a standard technique of inducing inflammation both *in vivo* [22, 23] and *in vitro* [24]. Depending on dosage, LPS treated animals display behavioural as well as cellular brain changes, predominately associated with glial activation [25, 26]. The current study investigated how acute systemic inflammation impacts upon *in vivo* cerebrovascular function and the status of the underlying NVU cells. This was investigated with a complementary set of *in vivo* neuroimaging measures in a rat model, paired with detailed characterisation of the cellular pathology of the NVU with immunohistochemistry methods.

## 2. Methods

### 2.1 Animals and pharmacological treatment

Female Hooded Lister rats (3-4 month old, 220g-320g) kept at a 12-hour light/dark cycle environment at a temperature of 22 °C with access to food and water *ad libitum* were housed in polycarbonate cages (*n*=3 per cage) in the Biological Services Unit at the University of Sheffield. Animals were fed conventional laboratory rat food. Sixteen animals were assigned to one of two groups (control *n*= 8 or LPS, *n*=8). Haemodynamic data were concurrently acquired in all treatment groups at two time intervals (4 and 6 hours) after injection, to characterise the development of the acutely induced LPS inflammatory response.

Each animal received an intraperitoneal injection based on condition. Control animals were administered a saline vehicle (1ml/kg), LPS animals received a dose of 2mg/kg LPS-EB (lipopolysaccharide from E.coli, 0111:B4 stain-TLR4 ligand, InvivoGen, Europe) dissolved in endotoxin-free water (InvivoGen, Europe), following loss of consciousness from anaesthesia.

### 2.2 Surgical procedures

Details of surgical and experimental paradigms were similar to those reported in previous publications from this laboratory [27-29]. Briefly, rats were anaesthetised with an intraperitoneal injection (i.p) of urethane (1.25mg/kg in 25% solution), with additional doses of anaesthetic (0.1ml) administered if necessary. Choice of anaesthetic was determined by urethane’s suitability for invasive surgery as well as long-lasting stability, which is essential in experiments where data collection lasts several hours [30]. Anaesthetic depth was determined by means of hindpaw pinch-reflex testing. Animals were tracheotomised to allow artificial ventilation and regulation of respiratory parameters. A left femoral artery cannulation was performed for mean arterial blood pressure (MABP) monitoring and blood gas analysis. A left femoral vein cannulation was also performed to allow continuous administration of phenylephrine in order to maintain blood pressure within a healthy physiological range (100-110 mmHg) [31].

To enable haemodynamic recordings, the skull was exposed via a midline incision and a section overlaying the left somatosensory cortex (barrel cortex) was thinned to translucency with a dental drill. This section was located 1-4mm posterior and 4-8mm lateral to Bregma [32]. A thinned skull was typically 100-200 μm thick with the local vasculature clearly observable. Care was taken during thinning to ensure that the skull remained cool by frequently bathing the area with saline.

### 2.3 Physiological monitoring

Temperature was maintained at 37 °C (± 0.5 °C) throughout surgical and experimental procedures with the use of a homoeothermic blanket and rectal temperature probe (Harvard Apparatus, USA). Animals were artificially ventilated with room air using a small animal ventilator (SAR 830, CWE Inc, USA); the breathing rate of each animal was assessed and modified according to each individual animal’s blood gas measurements. Respiration rates of the animals ranged from 68 to 74 breaths per minute.

Blood pressure was monitored during the experiment with a pressure transducer (Wockhardt, UK, 50 units of heparin per mL). Arterial blood from the femoral artery was allowed to flow back from the cannula into a cartridge (iSTAT CG4+, Abbott Point of Care Inc., USA) and analysed for blood gas to ensure normoxia and normocapnia using a blood gas analyser (VetScan, iSTAT-1, Abaxis, USA). Physiological parameters were within normal ranges throughout the experiment (mean values: PO_2_ =80mmHg(±9.1) PCO_2_ =30.4mmHg(±3.7) SO_2_ =96%(±1.2)) Total volume of arterial blood extracted at one time did not surpass 95μL. Phenylephrine (Sigma, Aldrich) was administered into the left femoral vein using a syringe pump (Sp200i, World Precision Instruments Inc., USA) to counteract reduced blood pressure caused by anaesthesia. Dose was adjusted accordingly to the blood pressure of each individual animal but remained in the range of 0.2-0.8mg/hr.

### 2.4 Imaging

Cerebral blood flow (CBF) data were acquired through a laser speckle contrast imaging (LSCI) camera (Full field Laser Perfusion Imager (FLPI-2), Moor Instruments, UK) which was positioned above the thinned cranial window. Images were acquired at 25Hz at a spatial resolution of approximately 10μm/pixel. A 70 second baseline data acquisition period was acquired to obtain a measure of baseline blood flow.

The two-dimensional optical imaging spectroscopy (2D-OIS) technique was used to estimate changes in oxygenated (HbO_2_), deoxygenated (HbR) and total (HbT) haemoglobin concentration in the rat barrel cortex. This technique has been previously described in detail [33, 34]. OIS data was collected at a frame rate of 8Hz. The spectral analysis was based on the path length scaling algorithm (PLSA) [33], which uses a modified Beer-Lambert Law with a path length correction factor. In our analysis, baseline haemoglobin concentration in the tissue was estimated to be 104μm based on previous measurements [35] and oxygen saturation estimated to be 50% when breathing room air.

### 2.5 Stimulation paradigms

Stimulation of the whisker pad was delivered via two subdermal stainless steel needle electrodes (12mmx0.3mm, Natus neurology Incorporated, USA) directly inserted into the whisker pad which transmitted an electrical current (1.0 mA). This intensity has been shown to evoke a robust haemodynamic response without altering physiological factors such as blood pressure and heart rate. The whiskers were stimulated at one of six frequencies (1, 2, 5, 10, 20 & 40Hz), for two seconds with a stimulus pulse width of 0.3ms. The order of stimulation frequencies was pseudorandomised with 10 trials at each frequency and an inter-trial interval (ISI) of 25s. The electrical current is generated by an independent amplifier (Isolated Stimulator DS3, Digitimer Ltd., UK) which directly attaches to the electrodes. All stimulation paradigms were carried out 4 and 6 hours after saline/LPS administration.

### 2.6 Hypercapnia challenge

A hypercapnia challenge is used as a measure of vascular reactivity which is independent of neuronal activity changes. During hypercapnia, a 10% concentration of carbon dioxide in medical air (9L medical air, 1L CO_2_) was administered to the air supply tube of the ventilator. Thirty-second long challenges were repeated four times at intervals of 210 seconds in the absence of whisker stimulation. An interval of 210 seconds ensured that the animal’s physiological parameters returned to baseline levels before delivering the next challenge. These challenges were performed following the two stimulation paradigms.

### 2.7 Perfusion

Rats (*n*=11) were transcardially perfused eight hours after LPS/saline administration (following *in vivo* data collection) with saline (0.9% warmed to 37 °C) with the addition of heparin (0.1ml/500ml) to exsanguinate the vessels and subsequently fixed in 4% paraformaldehyde (PFA) 01.M pH 7.4 in PBS. Saline and fixative were administered through a pump (Masterflex L/S, Cole-Parmer Instrument Company, UK) at a rate of 34ml/hr. Brains were stored in PFA overnight at 4 °C and subdissected in four regions and embedded in paraffin wax. Serial sections (5μm) were cut from the paraffin-embedded tissue.

### 2.8 Immunohistochemistry

Immunohistochemistry was performed using a standard avidin-biotin complex-horse radish peroxidase (ABC-HRP) method, and visualised with diaminobenzidine ([DAB], Vector Laboratories, UK). A summary of utilised primary antibodies and their conditions of use are shown in Table 1. Isotype and no primary antibody controls were included in every run and no specific immunoreactivity was observed.

**Table 1.** Antibody sources and experimental conditions

| *Antibody* | *Isotype* | *Dilution (time, temp)* | *Antigen retrieval* | *Supplier* |
| --- | --- | --- | --- | --- |
| Anti-GFAP | Rabbit IgG | 1:1000 (1 hr RT) | MW 10 min, pH9 | Dako, UK |
| Anti-IBA1 | Rabbit IgG | 1:400 (1 hr RT) | MW 10 min, pH9 | Abcam, UK |
| ICAM-1 | Goat IgG | 1:400 (1 hr RT) | MW 10 min, TSC pH 6.5 | R&D, UK |

### 2.9 Data Processing and Analysis

#### 2.9.1 In vivo imaging data

Data were processed in Matlab (2016a) using custom written code. Imaging data were spatially smoothed and then analysed using SPM [36], with regions of interest (ROIs) selected from a thresholded activation map and expected to include contributions from arterial, venous and parenchymal (capillary bed) compartments. Two animals, one from each group, were excluded from final analysis due to the absence of well-localised stimulus-evoked responses, thereby a total *n*=14 was used for final analyses (*n*=7 for each group). ROI size was consistent across animals, for both LSCI (*F*(1,13) = 0.06, *p*=.811) and OIS (*F*(1,13)=1.54 *p*=.239) data. LSCI and OIS time series of haemodynamic changes for each stimulation trials were then extracted from the ROI. LSCI time series were down-sampled to 5Hz (from 25Hz). Data from each stimulation trial were extracted and divided by the pre-stimulus baseline period (10s), to yield a measure of percentage change (fractional changes) in CBF, HbO_2_, HbR and HbT. Time series were averaged across trials according to stimulation condition. Area under the curve (AUC) and maxima for each response were calculated.

Statistical comparisons between groups using maxima and AUC response values were performed using multivariate ANOVAs (MANOVAs) for each time point (4 and 6 hours). A *p* value below .05 was considered to be a significant effect. Additional independent sample t-tests were conducted to further probe differences between experimental conditions. All statistical analyses were conducted using SPSS 23.

#### 2.9.2 Immunohistochemistry data

All images were taken from the contralateral side of the thinned cranial window (right side), to ensure inflammatory effects were not surgery (thinned cranial window) dependent. Images were taken from the somatosensory cortex (SS) and hippocampal subfields of the dentate gyrus (DG), CA1 and CA3. In the SS cortex three adjacent belt transects from the outer cortex through to the white matter border were taken for each animal at x20 magnification (Nikon microscope). SS area coordinates for captured imaged were taken between -.40 to -1.80 from Bregma (B) [37]. In the hippocampus, random field images were taken for each animal in the DG, CA1 and CA3 subfields. Hippocampus area coordinates for captured images were between - 3.30 to -5.30 from B [37]. All slides were imaged blind in a randomised order. Percentage GFAP, IBA-1 and ICAM-1 area immunoreactivity was quantified using Analysis^D^ software. All slides were analysed blind in a randomised order. One saline animal was excluded from analysis due to infection. Statistical analyses (independent sample t-tests) were performed using SPSS 23. A *p* value below .05 was deemed significant.

## 3. Results

### 3.1 Acute LPS treatment does not alter baseline CBF

To assess any effects of treatment upon baseline CBF the average perfusion value across a 30s period prior to the onset of stimulation at the start and end of the experimental protocol was calculated. A one-way ANOVA on CBF baselines at 4 hours (*F*(1,13)=0.10 *p*=.754) and at 6 hours (*F*(1,13)=0.59, *p*=.458) post treatment found no difference between groups.

### 3.2 Acute LPS treatment alters cerebrovascular responses to whisker stimulation

Multivariate analyses of variance (MANOVAs) were applied to HbO_2_, HbR, HbT and CBF response maxima or minima (HbR) values in order to determine significant effects of LPS administration at 4 hours and at 6 hours post treatment. All cases (*n*=14) were included in the analysis. At 6 hours post injection, LPS administration altered the profile of haemodynamic responses across the investigated stimulation frequency range with a significant interaction between stimulation frequency and (LPS or saline) treatment (*F*(20, 190) = 2.31, *p* =.002; Wilks’ Λ = .486). There were also significant univariate interaction effects for each haemodynamic measure as reported in Table 2. Post hoc analysis of individual stimulation frequencies using independent t-tests (Figure 1 A) was used to assess if any particular frequency was more salient in driving the above interaction effect. Results indicate a significant increase in HbO_2_ (*p*=.0496) and HbR (*p*=.022) response magnitude at 5Hz following LPS treatment. A representative haemodynamic response profile at a 5Hz stimulation frequency is plotted in Figure 1 B for each measure.

**Figure 1.**
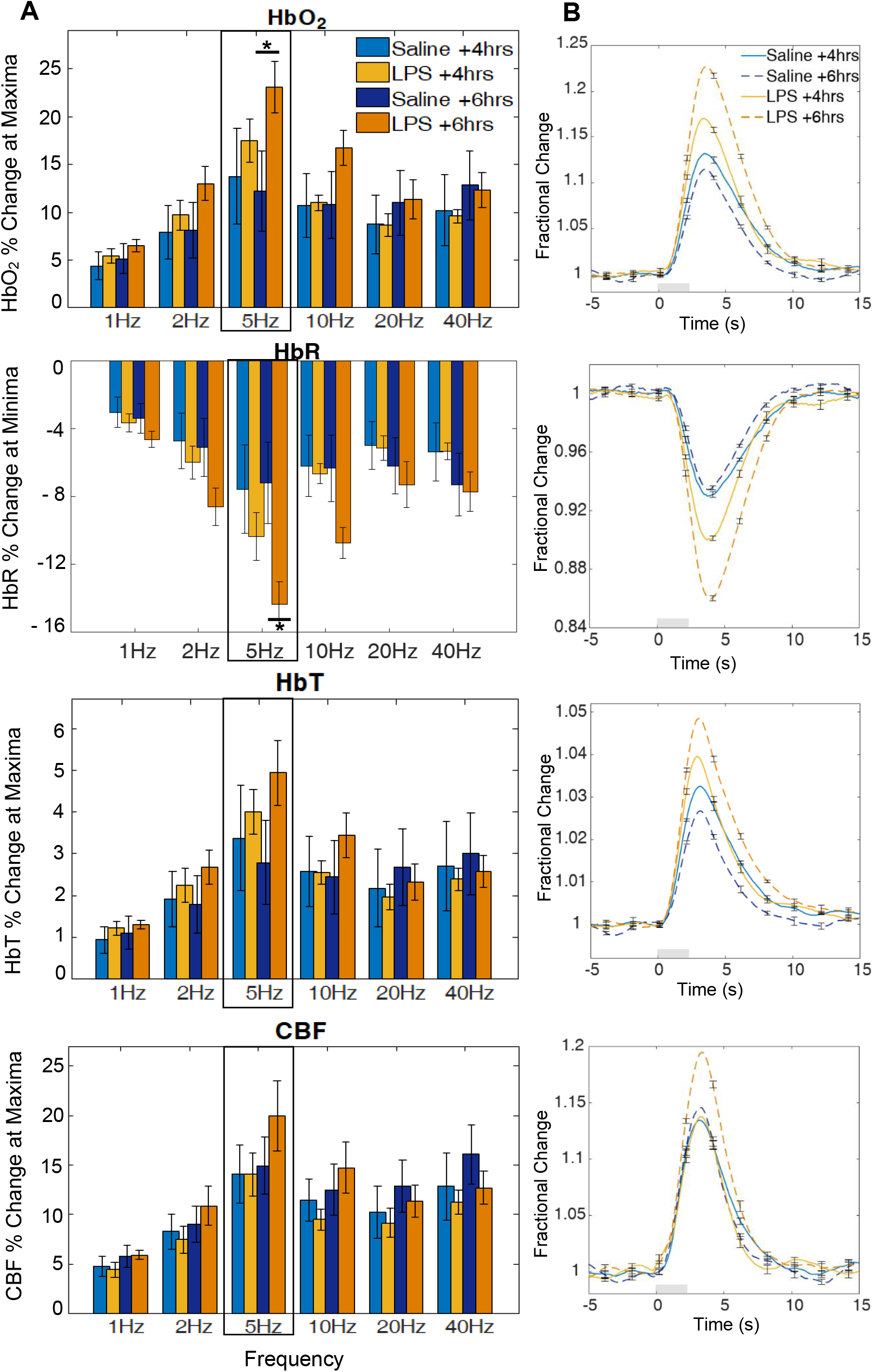
Haemodynamic response to 2s whisker stimulation at six frequencies at +4 and +6 hours after LPS/saline administration. **(A)** Bar charts show % change at maxima or minima (HbR). Post hoc analysis on stimulation frequencies reveal significant effects at 5Hz for HbO_2_ and HbR (*denotes significant differences at *p*<.05) between groups **(B)** Representative 5Hz time series showing mean fractional changes. Grey rectangle indicates stimulation onset/offset and error bars indicate standard error of the mean.

**Table 2.** Summary of univariate statistics for haemodynamic responses to a mixed frequency 2s stimulation paradigm.

| Time | Factor |  | *df* | *F* | *p* |
| --- | --- | --- | --- | --- | --- |
| +4 hours after injection | Frequency | HbO<sub>2</sub> | 5,60 | 15.19 | <.001 |
|  |  | HbR | 5,60 | 18.68 | <.001 |
|  |  | HbT | 5,60 | 11.80 | <.001 |
|  |  | CBF | 5,60 | 17.98 | <.001 |
|  | Frequency x Treatment | HbO<sub>2</sub> | 5,60 | 0.76 | =.581 |
|  |  | HbR | 5,60 | 1.45 | =.219 |
|  |  | HbT | 5,60 | 0.54 | =.743 |
|  |  | CBF | 5,60 | 0.23 | =.949 |
|  | Treatment | HbO<sub>2</sub> | 1,12 | 0.10 | =.758 |
|  |  | HbR | 1,12 | 0.24 | =.635 |
|  |  | HbT | 1,12 | 0.02 | =.892 |
|  |  | CBF | 1,12 | 0.15 | =.703 |
| +6 hours after injection | Frequency | HbO<sub>2</sub> | 5,60 | 21.45 | <.001 |
|  |  | HbR | 5,60 | 23.86 | <.001 |
|  |  | HbT | 5,60 | 15.25 | <.001 |
|  |  | CBF | 5,60 | 19.13 | <.001 |
|  | Frequency x Treatment | HbO | 5,60 | 6.48 | <.001 |
|  |  | HbR | 5,60 | 7.92 | <.001 |
|  |  | HbT | 5,60 | 4.75 | =.001 |
|  |  | CBF | 5,60 | 2.64 | =.032 |
|  | Treatment | HbO | 1,12 | 1.21 | =.294 |
|  |  | HbR | 1,12 | 2.36 | =.151 |
|  |  | HbT | 1,12 | 0.43 | =.524 |
|  |  | CBF | 1,12 | 0.07 | =.798 |

At 4 hours post injection, response magnitudes also appear altered, and especially for the 5Hz stimulation trials (Figure 1), but this is not supported by a MANOVA, with no significant interaction between stimulation frequency and treatment (*F*(20, 190) = 1.34, *p* =.156; Wilks’ Λ = .645).

At both time-points treatment condition by itself did not result in any significant effect on response magnitudes (4hrs: *F*(4, 9) = 0.33, *p* =.852; Wilks’ Λ = .872; 6hrs: *F*(4, 9) = 1.25, *p* =.357; Wilks’ Λ = .643). Thus although a consistent trend in increases in haemodynamic response was observed in LPS-treated animals, the interaction with stimulation frequency was key in driving significant differences between groups.

At both time points there was a significant effect of stimulation frequency on haemodynamic response magnitude (4hrs: *F*(20, 190) = 7.042, *p* <.001; Wilks’ Λ = .159; 6hrs: *F*(20, 190) = 6.41, *p* <.001; Wilks’ Λ = .181 [Figure 4.1 A]), indicating the range of stimulus inputs was effective in driving responses over a dynamic range, with significant effects for each haemodynamic response measure (Table 2).

### 3.3 LPS administration does not alter haemodynamic responses to hypercapnia

Analysis of variance (MANOVA) were used to determine significant effects of treatment on haemodynamic response magnitude (for HbO_2_, HbR, HbT and CBF variables) to a 30s hypercapnia challenge at 4 hours and 6 hours post treatment. Maxima (or minima for HbR) values were used to quantify the response. Treatment had no significant effect on response magnitude at 4 hours (*F*(4,9)=1.42, *p*=.304; Wilks’ Λ = .613) or at 6 hours (*F*(4,9)=1.33, *p*=.331; Wilks’ Λ = .629) (Figure 2).

**Figure 2.**
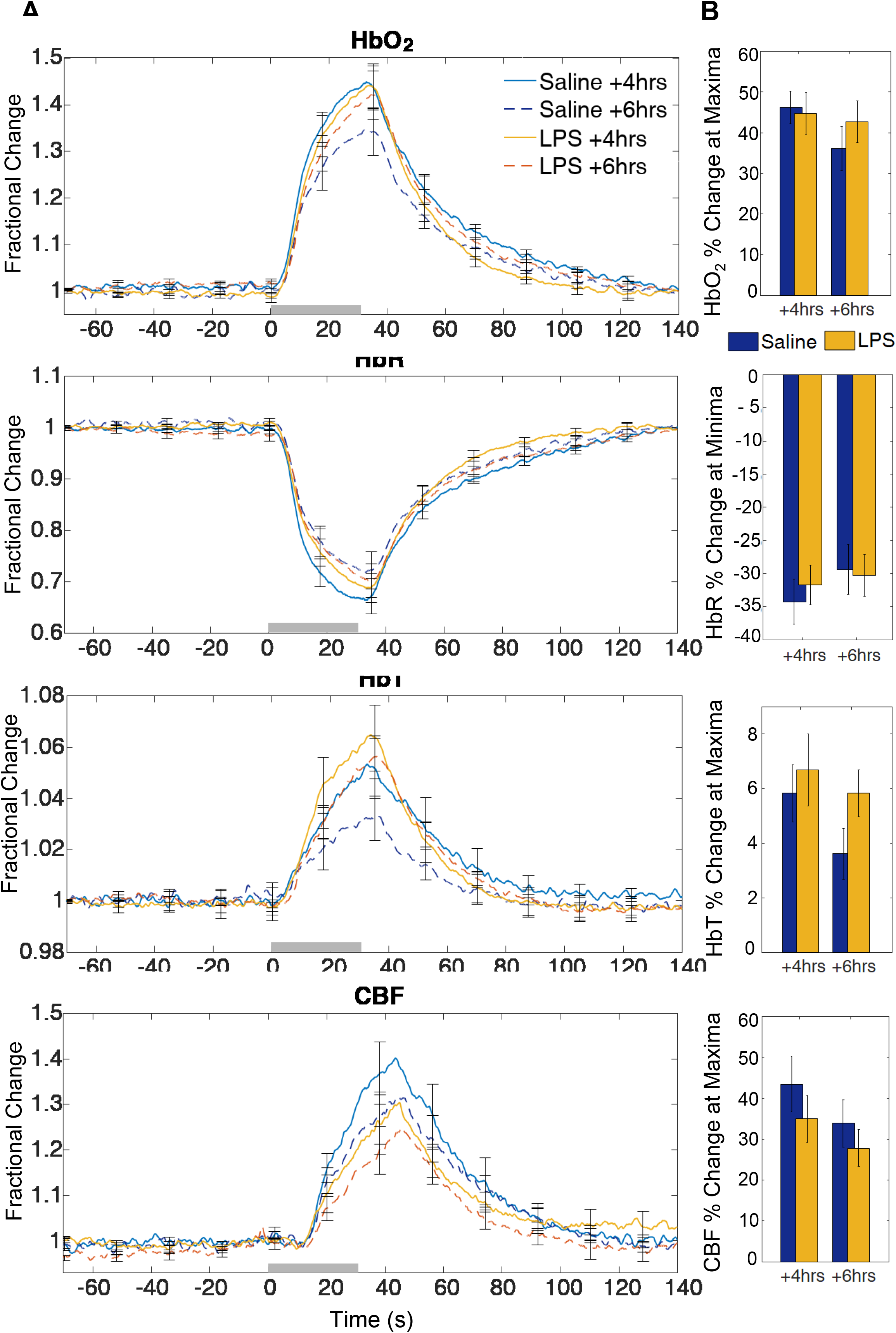
HbO_2_ HbR, HbT and CBF response to hypercapnia at +4hrs and +6hrs of LPS/saline administration. **(A)** Time series show mean fractional changes and **(B)** bar charts show percentage change at maxima or minima. Grey rectangle indicates the 30s CO2 challenge onset/offset. Error bars represent standard error of the mean.

### 3.4 LPS treated animals show a change in the cerebral metabolic rate of oxygen consumption (CMRO_2_) following treatment

The method utilised to estimate CMRO_2_ was as described in [28, 38]. Briefly the CMRO_2_ estimate was calculated from HbR, HbT and CBF values generated from OIS and LSCI data. CMRO_2_ in the brain is directly linked to cellular energy consumption and neuronal activity [39], thus it can provide a measure to assess the neurovascular coupling relationship by assessing changes in oxygen delivery or oxygen metabolism. CMRO_2_ changes were estimated at both time points (4 hours and 6 hours) and for both saline and LPS treated animals (Figure 3). Response maxima and AUC were calculated and analysed with one-way ANOVA. Whilst CMRO_2_ is estimated to initially increase in saline and LPS treated animals, in both LPS conditions (4 hours and 6 hours) there was a substantial decrease below baseline after the initial increase. There was no significant difference in the response maxima between groups at 4 hours or 6 hours after LPS/saline treatment (4 hours: *F*(1,13)=3.5, *p*=.087; 6 hours: *F*(1,13)=2.5, *p*=.138). However a one-way ANOVA on AUC reveals a significant difference at 6 hours (*F*(1,13)=6.16, *p*=.029) between treatment groups. No AUC significant difference was found at 4 hours (*F*(1,13)=2.05, *p*=.178).

**Figure 3.**
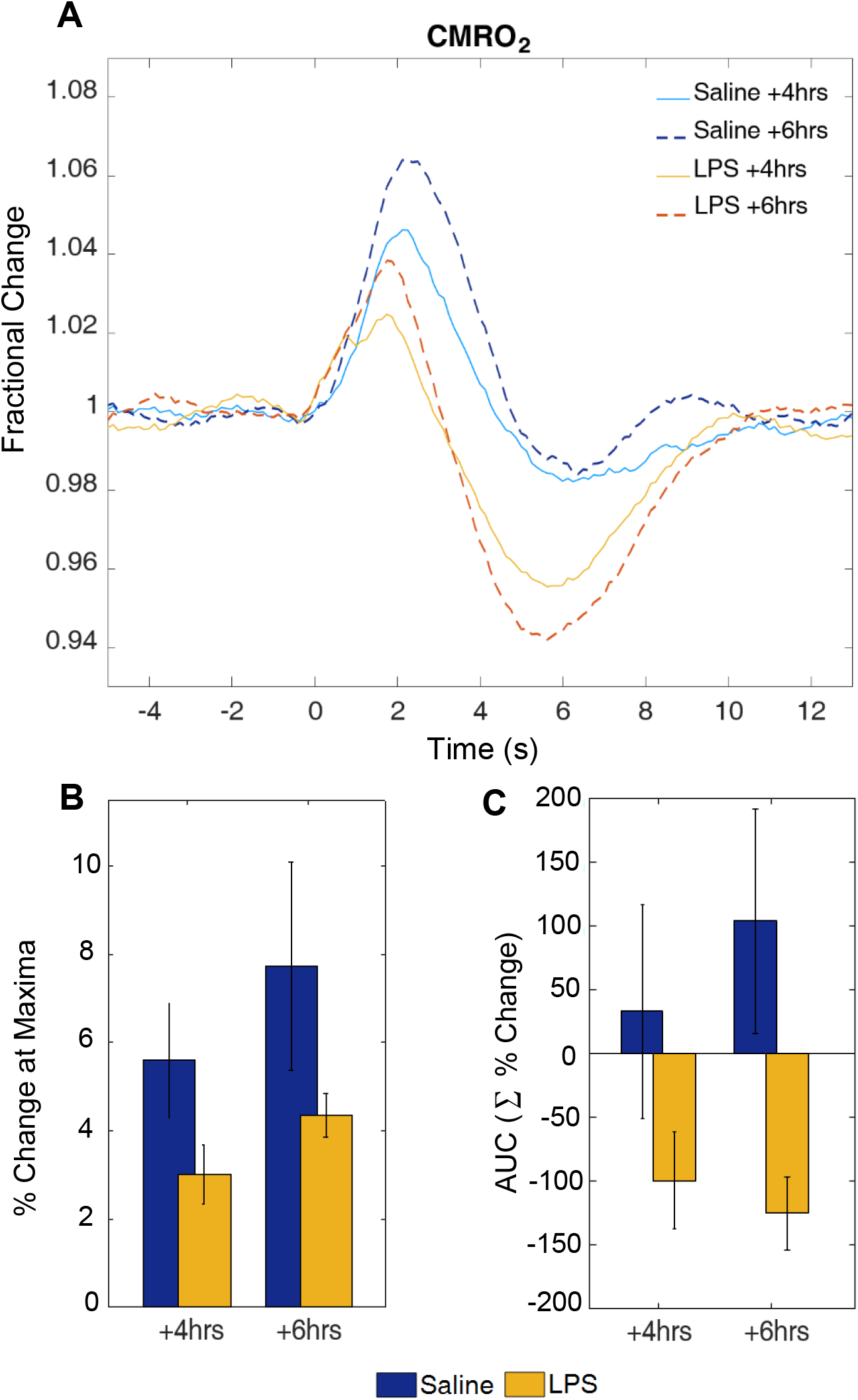
CMRO_2_ estimation at +4hrs and +6hrs of LPS/saline administration. **(A)** Time series of estimated CMRO_2_ changes following whisker stimulation, **(B)** Bar chart showing percentage change at maxima and **(C)** Bar chart showing change in area under the curve (AUC, units are summed percentage change).

### 3.5 Acute LPS treatment induces astrogliosis and microgliosis

GFAP immunolabelled the cell body and immediate processes of astrocytes in both cohorts (Figure 4 A-D). Hypertrophic astrocytes were observed in LPS cases, indicative of a mild to moderate astrogliosis phenotype. In LPS treated cohorts stronger immunoreactivity was observed around visible vessels compared to controls in both SS and hippocampal regions. In saline treated animals GFAP expression appeared to be predominately parenchymal (Supplementary Figure 1 A-D). Levels of GFAP expression, assessed as percentage area immunoreactivity showed a significant 74% increase in LPS cases in the SS cortex (*t*(8)= -4.15, *p*=.003) (Figure 4 I). Within the hippocampal subfields immunoreactivity was similar to control cases, no significant differences were found between LPS and saline treated groups (DG, *t*(8)= - 1.4, *p*=.200 (12% increase) (Figure 4 J), CA1, *t*(8)= -1.94, *p*=.089 [38% increase] or CA3, *t*(8)= - .34, *p*=.744 [6% increase]).

**Figure 4.**
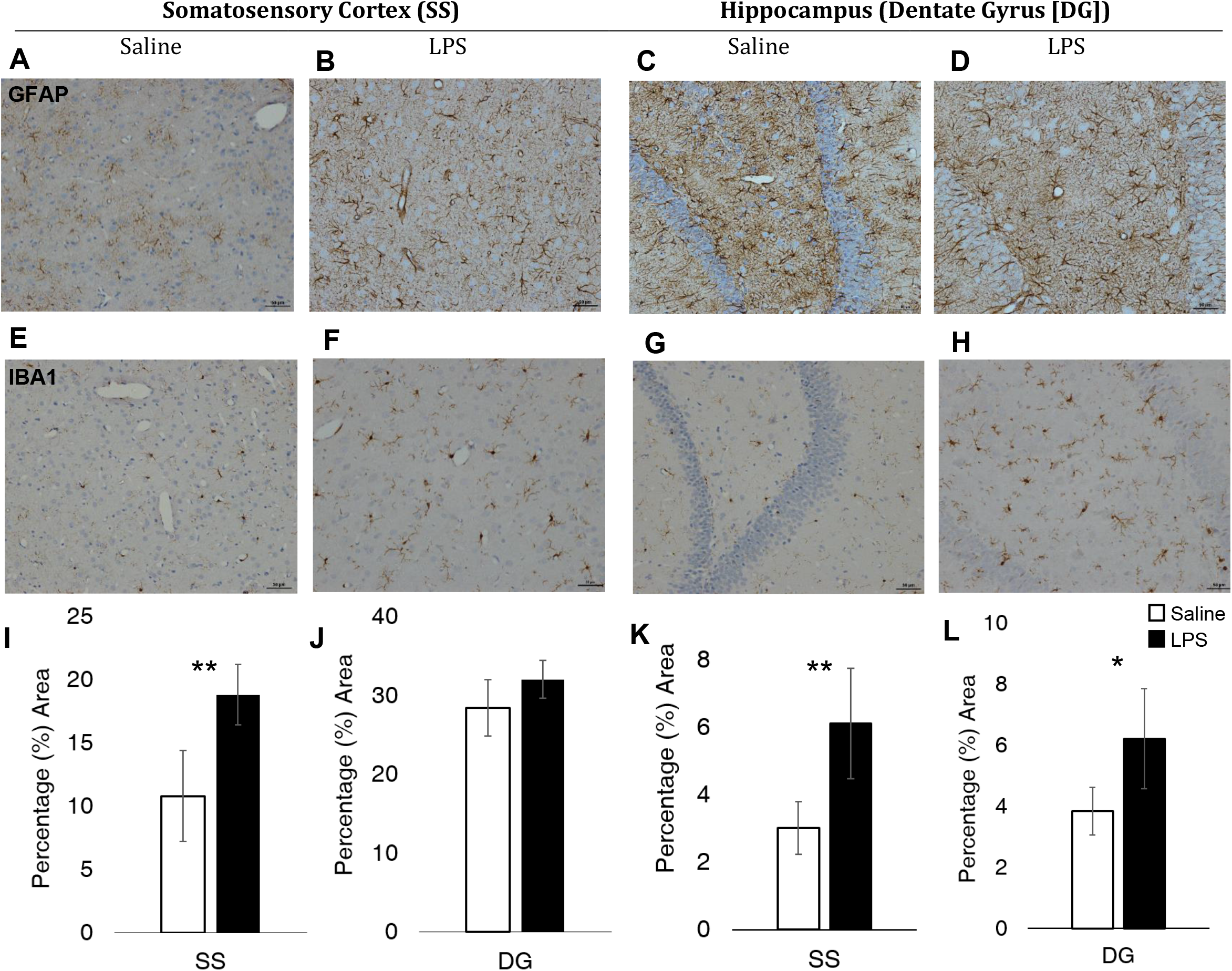
LPS-treatment changes in glial pathology. **(A-D)** GFAP immunolabelling of astrocyte cell body and immediate processes **(E-H)** IBA1 immunolabelling of microglia cell body and ramified processes. **(I)** GFAP SS percentage area immunoreactivity was significantly increased in LPS treated animals (74% increase, *p*=.003) but **(J)** was not significantly different in DG (*p*=.200). IBA1 SS **(K)** and DG **(L)** percentage area immunoreactivity were significantly increased in LPS treated animals (SS: 109% increase, *p*= .005; DG: 72% increase *p*=.036). Scale bar represents 50μm for **A-H.** ** denotes *p*<.010 * denotes *p*<.05.

IBA1 immunolabelled the cell body and ramified processes of microglia in both treatment groups (Figure 4 E-H). Similarly to GFAP expression, LPS IBA1^+^ microglia showed stronger immunoreactivity around vessels whereas, control cases displayed a predominant parenchymal distribution (Supplementary Figure 1 E-H). LPS-treated animals displayed a hypertrophic profile, indicative of a reactive microglial phenotype with a larger, more rounded cell body (Figure 4 F & H). Microglial clustering was observed along vessels (Supplementary Figure 2). Levels of IBA1 expression, assessed as percentage area immunoreactivity showed a significant 109% increase in LPS treated animals in SS cortex (*t*(8)= -3.84, *p*=.005) (Figure 4 K) with more intense immunolabelling of both the cell body and extending processes. In the hippocampus IBA1^+^ microglia immunoreactivity for LPS treated was significantly increased in all examined subfields (DG: *t*(8)= -2.52, *p*=.036 [72% increase] (Figure 4 L), CA1: *t*(8)= -4.09 [119% increase], *p*=.003, CA3: *t*(8)= -4.3, *p*=.003 [153% increase]).

### 3.6 Acute LPS treatment increased expression of ICAM-1 on the endothelial luminal surface and microglia processes

ICAM-1 immunolabelled vessels (Figure 5 A-D) in both cohort groups although LPS treated animals showed more intense immunoreactivity compared to controls. Furthermore, immunolabelling of microglia was a feature of all LPS cases but was less intense than vessel labelling. Percentage area assessment revealed a significant increase in ICAM-1 expression in LPS cases with a 299% area increase in the SS cortex (*t*(8)= -6.5, p<.001) (Figure 7 E) and in all hippocampal subfields compared to controls (DG: *t*(8)= -5.47, *p*=.001 [236% increase], (Figure 5 F) CA1: *t*(8)= -4.73, *p*=.001 [108% increase], CA3: *t*(8)= -4.69, *p*=.002 [186% increase]).

**Figure 5.**
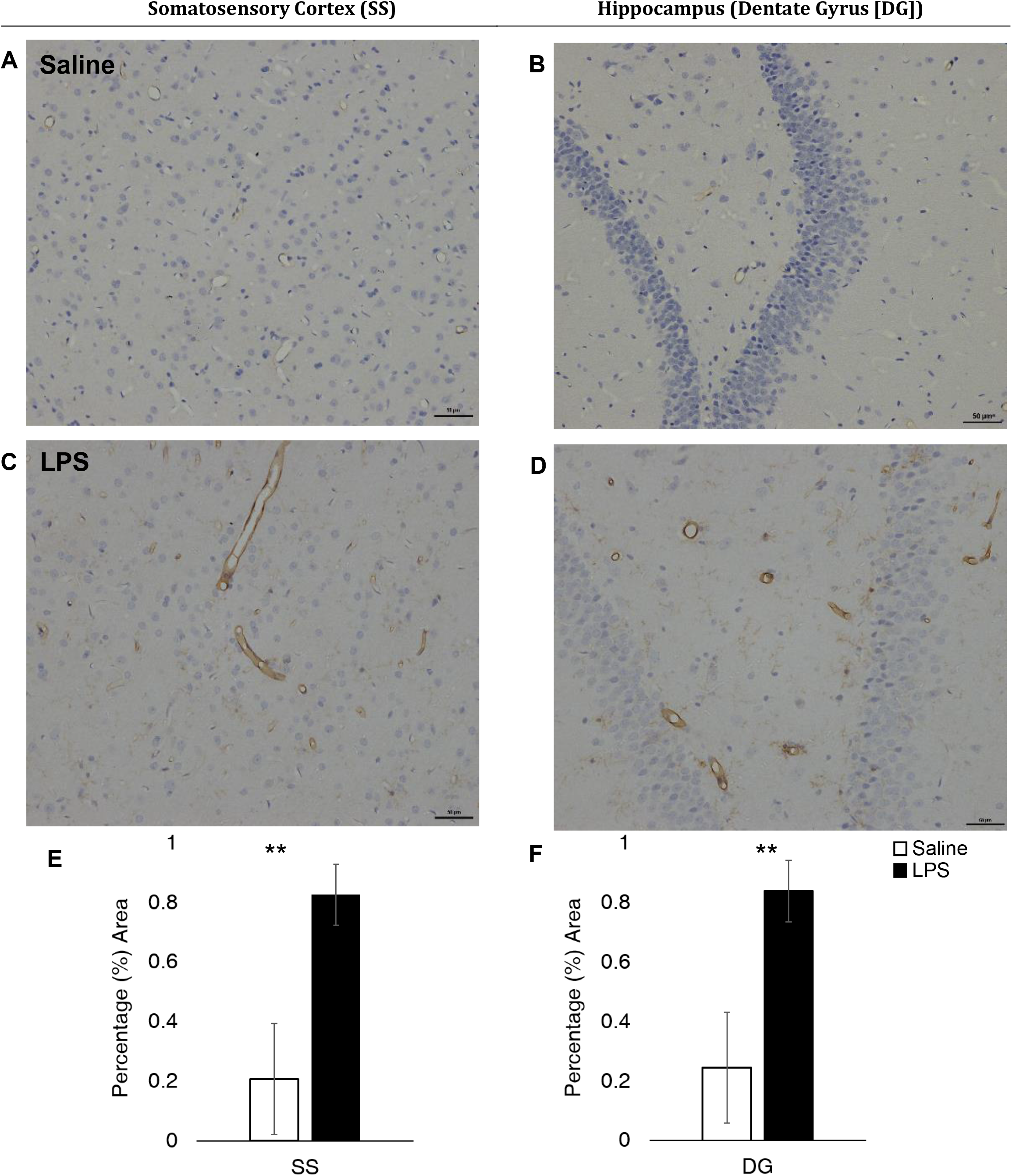
LPS-treatment changes in ICAM-1 expression. **(A-D)** ICAM-1 immunolabelled vessels and **(C-D)** microglia processes. LPS-treatment led to increased percentage area ICAM-1 immunoreactivity in the SS cortex **(E)** (299% increase, *p*<.001) and in the DG (F) (236% increase, *p*=.001). Scale bar represents 50μm for **A-D.** ** denotes p<.002.

## 4. Discussion

In this study we present evidence of an alteration of cerebrovascular function in animals acutely treated with LPS, demonstrated by changes in haemodynamic responses to a range of stimulus inputs. These findings have implications for the interpretation of neuroimaging data acquired in clinical and healthy cohorts where systemic inflammation may be present because physiological properties which underlie neuroimaging signals (CBF, CBV and CMRO_2_) acquired in humans may be altered. Moreover, characterisation of NVU cellular changes via immunohistochemistry showed morphological changes in astrocytes and microglia, alongside a marked increase in ICAM-1 expression, indicating ECs activation.

LPS administration significantly altered cerebrovascular function. These effects became evident at 6 hours after LPS administration, although similar (non-significant) trends could be observed at the 4 hour time point. It is difficult to assess why significant effects of altered cerebrovascular function are primarily driven by a specific stimulation frequency, although stimulation dependent effects have been previously reported in the literature [40, 41]. Our results suggest the 5Hz stimulation frequency to be particularly salient at driving this effect, and it has been previously reported that in anaesthetised animals this frequency is most effective in driving somatosensory neurovascular responses [41]. These results underline the complexity of the neurovascular coupling relationships, and the importance of including a dynamic range of sensory stimulations in neurovascular research to better assess how the neurovascular system responds to different levels of input in health and under disease-related manipulations.

CMRO_2_ in the brain is considered to be an index of brain health and energy homeostasis [39] as well as being directly linked to cell energy and neuronal activity [42]. Estimation of CMRO_2_ can thus provide a measure to assess the neurovascular dynamics by assessing possible changes in oxygen delivery or oxidative metabolism. CMRO_2_ estimations suggest an alteration in how oxygen delivery is matched to metabolic demand; where the increase or decrease of CBF relative to oxygen consumption is altered by LPS treatment. This alteration may in turn reflect a change in neurovascular coupling but this claim requires validation with a direct measure of neuronal activity. Nevertheless altered CMRO_2_ estimates indicate a change in the relationship between the HbR, HbT and CBF components of the haemodynamic response under LPS treatment, from which the CMRO_2_ estimate is made. This would in turn predict that BOLD signals, which are also derived from HbR, HbT and CBF changes will be different under LPS treatment, therefore caution should be taken in interpreting neuroimaging signals acquired from subjects with systemic inflammation present.

GFAP^+^ astrocytes in LPS-treated animals were more prominent, displaying greater immunoreactivity of processes and cell body. Furthermore immunoreactivity was also localised around vessels, suggesting increased GFAP expression by astrocyte endfeet. This phenotype is suggestive of mild to moderate astrogliosis [14]. A non-significant difference between GFAP^+^ immunoreactivity in the hippocampus of LPS treated and control animals may be explained by the dense population of GFAP^+^ astrocytes in the hippocampus compared to cortical areas [43, 44]. Further quantification utilising other GFAP isoforms alongside additional astrocyte markers (such as ALDH1L1, nestin, vimentin, EAAT1 and EAAT2) may help to fully elucidate the astrocyte response triggered by an acute LPS challenge. Similarly, characterisation of microglia revealed increases in IBA1 expression and a shift from a ramified ‘resting’ morphology to a hypertrophic profile characterised by an amoeboid appearance as seen in reactive microglia [12, 45]. Furthermore IBA-1^+^ microglia in LPS cases were observed to cluster around present vessels, a response possibly driven by the increase in ICAM-1 expression of the inflamed endothelium, as induction of ICAM-1 has been shown to correlate with increased microglial activation in acute inflammation [15].

ICAM-1 has an important role in cell-to-cell adhesion interactions and has low expression in cerebral microvessels in normal physiological conditions [46]. Its upregulation on the luminal surface of ECs is increased in the presence of pro-inflammatory mediators and has been linked to blood brain barrier (BBB) permeability during acute inflammation [15, 47]. In this study, we demonstrate increased ICAM-1 expression on the luminal surface of ECs, as well as microglia processes in LPS treated animals. ECs have already been shown to change their phenotypes in support of various phases of the inflammatory process [48, 49]. Furthermore EC location, at the interface between brain and blood, as well as their expression of TLR-4 receptors [50] (directly activated by the LPS strain used), could pinpoint these cells as initiators of the inflammatory response. Alterations in EC function could then propagate to the rest of the NVU cells, activating astrocytes and microglia. In support of this, a recent paper developing a concurrent cell-type specific isolation method [51], reported gene expression changes (upregulation of pro-inflammatory cytokines, chemokines, cell adhesion molecules, including ICAM1) in vascular endothelia in response to a peripheral LPS challenge. These results indicate that vascular ECs may be central in the initiation and transmission of the LPS response from the periphery to the CNS via cytokines, chemokines and extracellular remodelling [51].

ECs have also been implicated as key players in mediating neurovascular coupling in health [52]. Retrograde dilation of pial vessels following sensory stimulation is blocked if EC signalling is interrupted and disruption of ECs at the pial surface leads to a significant attenuation of the haemodynamic response [52]. These findings could help explain the increases in haemodynamic measures observed in this study, where such changes could be mediated by increased activity of ECs at the pial surface. Lastly, as ECs are exposed to the bloodstream, both BBB penetrating and non-penetrating substances (such as LPS) could act upon these cells to mediate changes in neurovascular coupling [53]. Intravenous administration of an anti-inflammatory substance (rofecoxib) was shown to attenuate cortical haemodynamic responses to sensory stimulation [54]. Rofecoxib inhibits production of prostaglandins (PGs), which are present in the cerebral endothelium. Furthermore, ECs through production and release of PGs, are key modulators of vascular tone and thereby can readily influence the haemodynamic response. As such it is possible that administration of LPS could be exerting its inflammatory effects through ECs/PGs increasing haemodynamic responses to sensory stimulation. Further investigation relating EC involvement in mediating haemodynamic changes and neurovascular coupling in health will be key in understanding EC roles in systemic inflammation and their impact upon neurovascular function and neurovascular coupling.

CNS cells including microglia [55] and astrocytes [56] possess TLR-4 receptors and thus should be directly activated by an LPS challenge. However, lipopolysaccharides and pro-inflammatory molecules are large and thus should have limited BBB permeability [57]. Evidence from research radioactively labelling LPS [58, 59] has shown little LPS penetration of the BBB (0.025% of the administered dose) and this level of penetration was only present at doses of 3mg/kg or higher. As such, in our model, we do not anticipate extravasation of LPS into the parenchyma. This physical limitation suggests that the observable brain inflammatory response produced by a peripheral administration of LPS is most likely mediated and initiated by ECs (which also have TLR-4 receptors, as previously discussed) and may also be mediated by alternative routes of communication between the brain and the periphery as opposed to a direct effect on glial TLR-4 brain receptors (for a comprehensive review see Holmes, 2013 [57]). This could explain why studies comparing brain and peripheral inflammatory challenges such as the one by Montacute et al. [60] report similar levels of brain inflammation in both challenges.

There is a pressing need to understand the communication pathways between the peripheral and central immune systems in order to understand the role of inflammation in neurological diseases and ageing and for the development of effective interventions combating inflammation. Neurodegenerative disease such as AD are characterised by a chronic low-grade inflammatory response, therefore a limitation of this study is the acute administration of an inflammatory challenge. It is, however, important to remember that as chronic and acute inflammation are not completely separate processes and share some of the same mediators, including EC activation [48], NVU changes measured in either acute or chronic inflammatory states can inform upon the other. Thereby acute alterations in *in vivo* neurovascular function, which arise from NVU cellular changes may also underpin the neurovascular changes observed in a chronic low-grade inflammatory response. Ageing, is furthermore, a key factor in both AD pathology and in inflammation. As such future work should aim to extend this model to include a low-dose chronically treated group with different aged animals to maximise its relevance to human pathology and human ageing.

Whilst we find no evidence of leukocyte extravasation in LPS treated cases, astrocytes and microglia are highly likely contributing to the neuroinflammatory response by secreting an array of cytokines. Future work should thus aim to characterise the inflammatory profile of glial cells in response to acute systemic inflammation. Lastly, varying species of LPS elicit different cytokines profiles, thereby producing different classes of immune response *in vivo* [61]. This is an important consideration for studies utilising LPS as a model of inflammation. The LPS strain utilised in this study, due to its ultrapure nature, only activates the TLR-4 pathway and thus offers a robust way of investigating the inflammatory-driven neurovascular and NVU effects which are mediated by a specific pathway.

This study has implications for the understanding of how cerebrovascular function changes under an acute inflammatory response. Assessing neurovascular function across different stimulation frequencies enables a detailed assessment on short-latency neurovascular impulse-response function [29, 41]. This is of particular importance as human fMRI research studies rely on short-latency impulse response functions to estimate the haemodynamic response [62]. It is thus important to robustly assess haemodynamic responses in preclinical research to better relate these findings to human fMRI studies and analysis. Furthermore the findings from the current study are relevant to the application of fMRI in subjects or patients with a systemic inflammatory response, as they show that measures underlying fMRI signals (CBF, CMRO_2_ and CB V) may be altered in a systemic inflammatory state.

## 5. Conclusion

Our data shows that an acutely induced systemic inflammatory response is able to rapidly alter *in-vivo* cerebrovascular function. Furthermore, it is associated with marked immunoreactivity within the cellular constituents of the NVU. We suggest the inflammatory response to be initially triggered by ECs as these cells are directly exposed to the bloodstream and have been implicated in mediating neurovascular function in health. Functional changes in ECs may then initiate a cascade of activation which propagates to other NVU cells such as glia.

## List of abbreviations

2D-OIS: : Two-dimensional optical imaging spectroscopy
Aβ: : Amyloid-beta
AD: : Alzheimer’s disease
AQP4: : Aquaporin 4
BBB: : Blood brain barrier
BOLD: :Blood oxygen level dependence
CBF: : cerebral blood flow
DAB: : 3,3’-Diaminobenzidine
DG: : Dentate gyrus
fMRI: : functional magnetic resonance imaging
GFAP: : Glial fibrillary acidic protein
IBA1: : Ionized calcium binding adaptor molecule 1
ICAM-1: : intercellular adhesion molecule
IL-1: : Interleukin 1
LPS: : lipopolysaccharide
LSCI: : Laser speckle contrast imaging
NVU: : Neurovascular unit
VCAM-1: : vascular cellular adhesion molecule
PET: : Positron emission tomography
SS: : Somatosensory cortex
TLR-4: : Toll-like receptor 4
TNF-α: : Tumour necrosis factor alpha

## Declarations

### Ethics approval and consent to participate

The present study was approved by the UK Home Office under the Animals (Scientific Procedures) Act 1986 and the University of Sheffield Animal Welfare and Ethical Review Body (AWERB, local ethics committee). All procedures were conducted under a U.K. Home office licence and have been reported in accordance with the ARRIVE guidelines.

### Consent for publication

Not applicable

### Availability of data and material

The datasets used and/or analysed during the current study available from the corresponding author on reasonable request.

### Authors’ con tributions

GB, CM and JS designed the study. CM and JS supervised the project GB collected and analysed *in vivo* and histology data. GB wrote the manuscript. GB, CM and JS interpreted the data. KA-A helped with initial histology data optimisation. All authors reviewed and edited the manuscript. All authors read and approved the final manuscript.

## Acknowledgements

The authors would like to thank the Sheffield Institute for Translational Neuroscience (SITraN) histopathology technical team, in particular Lynne Baxter, for their work in preparation of immunohistological protocols and for providing technical assistance when required.

---

**Supplementary Figure 1. Assessment of astrogliosis and microgliosis across brain regions.** LPS treated cases displayed higher immunoreactivity on vessels **(C-D)** as well as a parenchymal distribution. Astrocyte distribution in control cases is more parenchymal **(A-B).** IBA-1+ microglia in control cases appear to be distributed primarily in the parenchyma **(E-H)** whereas microglia in LPS cases display parenchymal localisation and are also more prominent around the vasculature **(G-H).** Scale bar represents 50μm.

**Supplementary Figure 2. IBA-1 vessel clustering.** LPS treated cases displayed microglial clustering in cortex and hippocampus. Representative image from the CA1 hippocampal region. Scale bar represents 50μm.

